# Protein engineering via Bayesian optimization-guided evolutionary algorithm and robotic experiments

**DOI:** 10.1101/2022.08.11.503535

**Authors:** Ruyun Hu, Lihao Fu, Yongcan Chen, Junyu Chen, Yu Qiao, Tong Si

## Abstract

Protein engineering aims to find top functional sequences in a vast design space. For such an expensive “black-box” function optimization problem, Bayesian optimization is a principled sample-efficient approach, which is guided by a surrogate model of the objective function. Unfortunately, Bayesian optimization is computationally intractable with the vast search space. Even worse, it proposes sequences sequentially, making it incompatible with batched wet-lab measurement. Here, we report a scalable and batched method, Bayesian Optimization-guided EVOlutionary (BO-EVO) algorithm, to guide multiple rounds of robotic experiments to explore protein fitness landscapes of combinatorial mutagenesis libraries. We first examined various design specifications based on an empirical landscape of protein G domain B1. Then, BO-EVO was successfully generalized to another empirical landscape of an *Escherichia coli* kinase PhoQ, as well as simulated NK landscapes with up to moderate epistasis. This approach was then applied to guide robotic library creation and screening to engineer enzyme specificity of RhlA, a key biosynthetic enzyme for rhamnolipid biosurfactants. A 4.8-fold improvement in producing a target rhamnolipid congener was achieved after examining less than 1% of all possible mutants after 4 iterations. Overall, BO-EVO proves to be an efficient and general approach to guide combinatorial protein engineering without prior knowledge.

## INTRODUCTION

A protein fitness landscape describes the metaphoric, high-dimensional surface relating a select property (“fitness”) to amino acid sequences (1). Exploring this landscape via protein engineering is challenging for the following reasons: (i) The search space grows exponentially with the number of amino acid positions considered; (ii) Functional proteins are extremely rare, and high-performance sequences decrease exponentially with increasing fitness (2, 3); (iii) The fitness landscape is rugged attributing to epistasis (4); (iv) Experimental profiling is laborious, expensive, and slow. As a powerful protein engineering strategy, directed evolution performs multiple rounds of variant library creation and screening (1, 5), which often fixes one top mutation in each round. However, such greedy exploitation may be trapped in local optima, especially in a rugged fitness landscape due to epistasis among mutations.

Machine learning (ML) algorithms are increasingly applied to both model fitness landscape and guide protein engineering (6). To learn sequence-activity relationships from labelled data, supervised models have been created to predict various properties including thermostability (7), fluorescence (8), ligand-binding affinity (9), and catalytic performance (10). On the other hand, gene and protein sequences accumulate at an unprecedented speed in public databases (e.g., UniProt (11)). Unsupervised models, such as UniRep (12), TAPE (13), ESM-1v (14), and ProtT5-XL-U50 (15), have been developed to learn representation from vast unlabelled data and discover latent patterns in protein sequences. Although achieving proof-of-concept successes, the performance of current fitness models is impaired by data scarcity and bias (16). For example, the fitness levels of evolutionarily sampled sequences are generally in the medium range, and therefore the label diversity is limited in the low- and high-fitness ranges. Also, as functional assays are expensive and laborious, labelled data often cover a tiny fraction of the whole sequence space, resulting in a learned model with pathology outside the scope of training sets.

To guide protein engineering, both supervised and unsupervised ML models have been utilized to improve sample efficiency. For example, machine learning-assisted directed evolution (MLDE) first employed a zero-shot, physics-based model to calculate ΔΔG values of all possible mutants and filter out less-stable mutants. Then, an experimental set of 384 candidates was selected to train fitness models, which were applied to prioritize top candidates by *in silico* fitness evaluation (17). As a result, MLDE was 81-fold more frequent than greedy directed evolution to reach the global fitness optimal, when applied to a 4-site combinatorial landscape of protein G domain B1 (GB1). Furthermore, the evolutionary context-integrated neural network, ECNet, exploited homologous sequence information in a supervised way to predict high-order mutational effects, and successfully engineered TEM-1 β- lactamase with 2 to 6 mutations conferring up to 8-fold enhancement in ampicillin resistance (18). Moreover, utilizing learned representation from >20 million natural sequences by unsupervised models, the low-*N* protein engineering method required as few as 24 sequence-function data to train a supervised fitness model and exploited the model using simulated annealing. As a result, top mutants were successfully identified from reported mutant libraries of GFP and TEM-1 β-lactamase proteins (19). Notably, current ML-guided approaches seek to keep experimental batch size and iteration number as low as possible, confined by the physical limit of human researchers.

On the other hand, biofoundries accelerate design-build-test-learn cycles in biological engineering through physical and informatic automation (20–22). When applied to protein engineering, automation permits large-scale library creation and screening in a short time, so that new sequence-function data of sufficient amount and quality can be iteratively collected (23, 24), leading to continuous improvement of both model prediction and sequence design. However, the enhanced affordability and availability of robotics-generated data have not been well exploited in algorithm-guided protein engineering. To achieve efficient feedback between ML algorithms and robotic experiments, three modules are necessary for performing data acquisition, fitness modelling, and sequence proposal in iterations. The data acquisition module sends sequence queries to protein engineering robots and returns the corresponding fitness values. The fitness modelling module then performs model refinement on the newly acquired sequence-function data. Guided by the updated predictive models, the sequence proposal module generates new sequences for *in silico* evaluation, ideally accounting for both predicted fitness and model uncertainty. In this regard, Bayesian optimization or BO (25) is well suited for stepwise optimization on complex fitness landscapes, whereas model uncertainty can be explicitly considered to negotiate exploration and exploitation using a defined acquisition function. Indeed, BO has been successfully applied to engineer proteins (26), pathways (27, 28), and fermentation strains (29).

Two main customizations to BO are necessary to guide protein engineering on combinatorial fitness landscapes. First, whereas BO takes only one new query each time, a batch of mutant sequences can be readily created and evaluated in parallel by robotics. In this aspect, batched BO has been practiced to design biological sequences (30). There are two general categories of batching strategies: building batches iteratively (31) or using acquisition function over batches (32). Nevertheless, it remains elusive what the best strategy is to balance exploration and exploitation of batched BO. Second, BO locates the global optimum via brute-force enumeration over the entire search space. For combinatorial mutagenesis of proteins (N residues ~ a 20^N^ design space), the required computing resources for BO scales exponentially with sequence length. Therefore, heuristic methods (33, 34), such as evolutionary algorithms, are expected to restrict the BO search space. For example, genetic algorithms generate a batch of sequence candidates using random mutation and recombination, and top-performing mutants can be identified by fitness models. However, the use of evolutionary algorithms alone is often greedy and less exploratory, so it is desirable to integrate evolutionary and BO methods for the efficient and scalable study of combinatorial fitness landscapes.

Here, we developed a batched, BO-guided EVOlutionary algorithm (BO-EVO) for protein engineering via iterations of ML models and robotic experiments. Using an empirical GB1 dataset of 4-residue combinatorial fitness landscape (20^4^=160,000) (35), we explored and optimized many design specifications, including experimental budget, sequence representation, and iteration initialization, and identified surrogate model quality and moderate batch size as key considerations. The generalization capability of BO-EVO was evaluated using another empirical landscape of PhoQ (36) and a reformulated version of mathematical NK landscape (37), which reveals landscape ruggedness as a limiting factor. Excitingly, BO-EVO successfully guided iterative robotic engineering of RhlA mutants, achieving a 4.8-fold improvement in selective production of a rhamnolipid congener after only 4 iterations.

## MATERIAL AND METHODS

### BO and GPR methods

BO (25) is a derivative-free approach to optimize an objective function that is expensive to evaluate and lacks known structures like concavity or linearity. BO builds a surrogate model of the objective, estimates model uncertainty, and utilizes an acquisition function to negotiate exploration and exploitation and decide where to sample next. We used Gaussian process regression or GPR (38) to model the objective. A Gaussian process (GP) is a generalization of the Gaussian probability distribution. Whereas a probability distribution describes random variables which are scalars or vectors, a stochastic process governs the properties of functions. Leaving mathematical sophistication aside, one can loosely think of a function as a very long vector, each entry in the vector specifying the function value *f*(**x**)~Normal(μ(**x**),σ^2^(**x**)), at a particular input **x**. A GPR uses GP for regression and is fully described by a mean function *μ(**x**)* and a covariance function or kernel Σ(**x, x′**). The kernel is chosen so that points **x**, **x′** that are closer in the input space have a large positive correlation, encoding the belief that they should have more similar function values than points that are far apart. In this article, zero mean function prior was used, and a scaled radial basis function (RBF) kernel was applied, 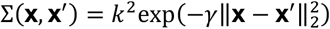, where *γ* can be interpreted as a similarity measure of the input space and *k* is related to prior uncertainty. The problem of learning in GP is exactly the problem of finding suitable properties for the kernel, specifically, determining the hyperparameter *k* and *γ*. We used a marginal log-likelihood as a loss function to estimate the kernel parameters using training sets and inferenced for a new query **x**_q_ according to the learned posterior to obtain the mean *μ(**x**_q_)* and variance σ^2^(**x**_q_). The readers are referred to (38) for more details. To accelerate the training and inference procedures, we used GPyTorch (39) with CUDA acceleration as a more efficient GP implementation than that offered by scikit-learn. Regarding training, maximum likelihood estimation was performed with Adam optimizer.

We used the upper confidence bound or UCB (40) as an acquisition function, which is to be optimistic in the face of uncertainty. And the Gaussian process upper confidence bound (41) was proposed as a Bayesian optimistic algorithm with provable cumulative regret bounds in the bandit setting. In the GP case, since the posterior at any arbitrary point **x** is a Gaussian, any quantile of the distribution of *f*(**x**) is computed with its corresponding value *β* as follows: *α_UCB_*(**x**) = *μ*(**x**) + *βσ*(**x**). The hyperparameter *β* was tuned to achieve optimal performance.

### Genetic algorithm and fitness-based sampling

Vanilla genetic algorithms are population-based. The algorithms generate a child population through recombination and mutation of a parent population and screen top candidate children for the next round of evolution. Instead, we only used random mutation on one parent to generate a child population. The per sequence mutation rate is set to introduce a single mutation in one sequence on average. One parent sequence is mutated into 6 to 10 child sequences suggested in Figure S3. For a subsequent round, the parent sequence is either set as the top child sequence screened or directly sampled from measured sequences following a fitness-based sampling strategy, where a higher fitness sequence is sampled with higher probability proportional to the exponential function value of its fitness.

### Protein fitness landscapes

To develop, validate and verify BO-EVO, diverse fitness landscapes were used. First, we developed BO-EVO on the empirical fitness landscape corresponding to a 4-site combinatorial mutagenesis library of GB1 (35). Then, we validated the algorithm on an additional empirical protein fitness landscape, PhoQ (36), and simulated NK landscapes reformulated from the original NK landscape (37). The fitness distributions are given in Figure S1. Finally, we verified the algorithm by guiding multiple rounds of robotic engineering to generate improved RhlA mutants. Details of the landscapes are given below.

#### GB1 landscape

GB1 is a 4-site (Val39, Asp40, Gly41, and Val54) combinatorial empirical fitness landscape of protein G domain B1 (PDB ID: 2GI9). The landscape consists of 20^4^ = 160,000 variants in theory, within which 149,361 variants were experimentally measured and the rest were computationally imputed. Variant fitness was determined by both stability (i.e., the fraction of folded proteins) and function (i.e., binding affinity to lgG-Fc) (35). Normalized by the global maximum minus the global minimum, the fitness value ranges from 0 to 1, with the WT fitness equals to ~0.1.

#### PhoQ landscape

Like GB1 landscape, PhoQ (UniProt ID: P23837) is also a 4-site (Ala284, Val285, Ser288, and Thr289) combinatorial mutagenesis landscape. Fitness reflects protein-protein interaction (PPI) between an *E. coli* kinase PhoQ and its cognate protein substrate PhoP (36). The experimental coverage of the fitness landscape is nearly complete (140,517 out of 160,000). The same fitness normalization strategy was applied as with GB1, and the WT fitness is about 0.02.

#### NK landscapes

The NK landscape, proposed by Kauffman and Weinberger (37), is a statistical model of combinatorial fitness landscapes that captures *epistasis* as the key characteristic. Several parameters are explained: *N* gives the length of the sequences; *K* gives the number of sites that interact with a select site; and thus, *K* represents the order of epistasis in the landscape; *V* gives the number of alternatives at each site; for proteins, *V* = 20 for classical amino acids. For any site *i*, the interacting set of sites is *v_i_* = {*v*_*i*,1_,*v*_*i*,2_,…,*v_i,k_*}. These are typically assigned uniformly and independently at random. The fitness of some sequence *X* of length *N* is given by 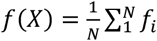, here *f_i_* is the “fitness contribution” associated with a site *i*. The fitness contribution is determined by the combination of *x_i_* and {*x_j_*}_*j*∈*v_i_*_. There are 20^K+1^ combinations of *K* + 1 amino acids and each of these combinations is assigned by selecting an independent random value from the uniform distribution on (0,1). This constitutes the “fitness table” for the site *i*. There is a different, independently generated table for each of the *N* sites. Finally, the fitness is normalized by global maximum minus global minimum, resulting in the fitness value ranging from 0 to 1, and we set the fitness of WT sequence as 0.1 (similarly hereinafter). We name this NK landscape “Original_NK”.

Accordingly, the fitness of the “Original_NK” landscape is normally distributed, whereas that of empirical protein fitness landscapes obeys exponential distribution (see Figure S1). To mimic empirical fitness landscapes, the fitness contribution was assigned by sampling from an exponential distribution instead of the uniform distribution. This resulted in an exponential distribution of the high fitness values. We named the resultant landscape “Exp_NK”.

As depicted by the above equation, *f_i_* contributes equally to sequence fitness *f*(*X*), ignoring the different contributions of each site. However, only a small number of key residues exert substantial impacts on a target function. Thus, we proposed a weighted average of *f_i_* for *f*(*X*), with the weight sampled from a Zipf distribution. We named the landscape “Exp_NK_Zipf’.

Furthermore, functional proteins cover only a tiny fraction of the whole sequence space (2). Hence, we simply set the low-fitness variants in the “Exp_NK_Zipf’ landscape to “non-functional” with a cut-off threshold of 0.3 and created a black hole-filled, exponentially distributed NK landscape. We named the landscape “Exp_NK_Zipf_hole”.

#### Ruggedness-to-slope ratio, r/s

Various measures of epistasis in the literature appear to be equivalent (42), and here we used the ruggedness-to-slope ratio (43), *r/s*, to measure landscape ruggedness. This ratio measures how well the landscape can be described by a linear model, which represents the purely additive (non-epistatic) limit. It is obtained by fitting a multidimensional linear model to the empirical fitness landscape through a least-square fit. The linear model is 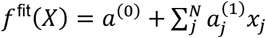, where the parameters *a*^(0)^ and the 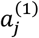 s are fitted. The mean slope is 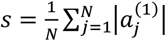 and the roughness is defined by 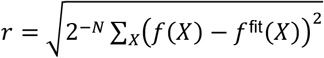. The higher *r/s*, the higher the deviation from the linear model and the greater the level of epistasis present in the landscape. The *r/s* ratio of several landscapes are plotted in Figure S2.

### Fitness landscape exploration algorithms

In addition to BO-EVO, two baseline exploratory algorithms (random solver and evolutionary solver), as well as a batched enumerated BO, are implemented for performance comparison. Several key parameters are listed in Table S2.

#### Random

We used Random solver as a weak baseline solver. Random solver generates new sequences by random mutation of parent sequences with a per-site mutation rate of 1/N, where N is the sequence length. And the parent sequences are randomly selected from the previously measured batches. M candidate sequences are generated, and a batch of B sequences are randomly selected from the candidates for measurement.

#### AdaLead

It is an evolutionary greedy solver proposed by Sinai et al. (44). The algorithm selects a set of parent sequences such that the fitness is within (1 – *κ*) of the maximum fitness measured in the previous batches. These parents are then iteratively recombined and mutated. Evaluated by surrogate models, children sequences showing fitness improvement over the corresponding parent are added to a set of candidates M. And finally, all candidates are ranked according to predicted fitness by the surrogate model, and the top B sequences are proposed for measurement.

#### BO

It is a batched enumerative Bayesian optimization algorithm, which generates batch sequences by exhaustively evaluating the acquisition function and prioritizing all the sequences in the whole design space. The top B new sequences are proposed as a new batch for measurement. As with BO-EVO, the surrogate model is GPR, and the acquisition function is UCB.

### Simulation setup and evaluation metrics for landscape exploration algorithm

#### Starting sequence

To develop algorithms for practical use, we randomly selected a batch of starting sequences with similar fitness levels as WT for each landscape. For the GB1 landscape, for example, the fitness values of starting sequences are close to 0.1, which is the WT fitness. The starting sequences were selected as follows:

1. Filtering the library with a fitness value interval (0,0.2);
2. Splitting the fitness interval (0,0.2) uniformly into 5 bins, and grouping the filtered variants into 5 groups according to their fitness;
3. Grouping the variants in each group into subsets according to its hamming distance to the global optima;
4. Sampling one variant from each subset as a starting sequence, randomly.

Considering that the exploration algorithms use random procedures to generate new sequences, five random seeds, and hence five simulations, were applied for each starting sequence to eliminate random factors. Consequently, a total number of (#starting_sequence×5) simulations are conducted for each landscape.

*Success ratio* is the fraction of simulations in which algorithms reach the global maximum or a target fitness level. *Cumulative maximum fitness* is the cumulative maximum of measured fitness sampled until each round in each simulation. *Maximum fitness* is the maximum of measured fitness sampled in the iterations in each simulation, whereas *mean fitness* is the mean of that.

### Protein sequence encoding strategies

Model architectures and datasets used for the pre-trained representation models are summarized in Table S3.

#### Onehot

Amino acids are encoded as one-hot vectors. The dimension is C=20 for classical amino acids. Then the one-hot vectors are concatenated into a long vector of dimension N×C for the entire sequence of length N (similarly hereinafter).

#### Georgiev

The physicochemical properties of amino acids, such as hydrophobicity, volume, and mutability, are considered by constructing a low dimensional vector of over 500 amino acid indices from the AAIndex database (45) by the primary component analysis (PCA) approach (46). The manually curated indices describe amino acid properties. The vector dimension for each residue is C=19.

#### UniRep

It is a contextual embedding learned by a 1,900-dimensional single hidden layer multiplicative LSTM model, which was trained on the UniRef50 dataset (47). The pre-trained model used is “babbler-1900”, and hence the dimension of the vector for each residue is C=1,900 (12).

#### TAPE

The feature is a contextual embedding learned by the transformer encoder, in the way of BERT (48). TAPE consists of 768-dimensional 12 hidden layers, and was trained on the dataset collected from Pfam database (49). The pre-trained model used is “bert-base”, and the features at the final hidden layer were extracted for an input sequence. Thus the dimension of the vector for each residue is C=768 (13).

#### ESM-1v

The feature is also a transformer-style embedding, which is specialized for mutation effects prediction. The BERT model, which consists of 1,280-dimensional 33 hidden layers, was trained on the dataset collected from UniRef90 dataset (47). The pre-trained model is “esm-1v”, and the features at the final hidden layer were extracted for an input sequence. Thus the dimension of the vector for each residue is C=1,280 (14).

#### ProtT5_XL_U50

Unlike the previous language models, ProtT5-XL-U50 uses an encoder and a decoder, T5 model (50). The model consists of 1,024-dimensional 24 hidden layers, containing 3 billion parameters. It was trained on the dataset collected from BFD dataset (51, 52) and fine-tuned on the dataset collected from UniRef50 dataset (47). The pre-trained model is “prot_t5_xl_uniref50”, and the features at the final hidden layer were extracted for an input sequence. Thus the dimension of the vector for each residue is C=1,024 (15).

### DNA, strains, and cultivation conditions

The cloning strategy and primer list can be found in Table S5 and Table S6, respectively. Enzymes used for molecular biology, including Q5 High Fidelity DNA Polymerase, restriction enzymes, and ligases were from New England Biolabs (NEB, Ipswich, MA). Spin Plasmid Mini-prep Kits (Axygen) were utilized to isolate plasmid DNA from *E. coli*. PCR, digestion, and ligation products were purified by PCR Purification and Gel Extraction Kits (Axygen). Gibson assembly was performed using NEBuilder HiFi DNA Assembly Master Mix (NEB) following the manufacturer’s instructions. Golden Gate assembly was set up with 25 ng of each plasmid, 0.15 μL of BsaI-HF or BsmBI, 0.15 μL of T4 ligase, 0.5 μL of 10×T4 ligase buffer, and water in a total reaction volume of 5 μL. The thermocycling program included an initial 5 min incubation at 37°C, 30 cycles of alternating 5 min incubation at 37°C, and 5 min incubation at 16°C, followed by infinite incubation at 16°C until *E. coli* transformation. Sanger Sequencing was performed at Sangon biotech (Shanghai, China).

*E. coli* DH5α (AlpaLife, Shenzhen, China) was used for plasmid amplification and library construction. *E. coli* BL21(DE3) (AlpaLife, Shenzhen, China) was used for rhamnolipid production. *E. coli* strains were statically cultured at 37°C on agar or aerobically cultivated at 30°C or 37°C and 220 rpm in LB media supplemented with 50 μg mL^-1^ kanamycin (LB-Kan). Transformation of chemically competent *E. coli* was performed via heat shock at 42°C. For inducible mono-RL production, isopropyl β-D-1-thiogalactopyranoside (IPTG) was supplemented at a final concentration of 1 mM. Chemicals were purchased from Sigma-Aldrich (Schnelldorf, Germany) or Sangon biotech (Shanghai, China).

### Robotic processes for molecular biology, microbiology, and MS analysis

#### Construction of mutant plasmid libraries

Site-directed mutagenesis (SDM) and site-saturation mutagenesis (SSM) were performed using Gibson assembly of two PCR products generated using primers containing a single mutation or the NNK degenerative codons (Figure S5 and Table S6) as we previously reported (23). Briefly, PCR products were treated with 1 μL of DpnI at 37°C for 1 h to remove any carryover of plasmid templates. Purified PCR products were then used for Gibson Assembly and *E. coli* DH5α transformation. After 16 h incubation at 37°C on LB-Kan agar, independent clones were randomly picked using QPix colony picker into 1 mL of LB-Kan in 96-deep well plates. After 20 h cultivation at 37°C, half of the resulting cultures were subjected to Sanger sequencing. For SSM libraries, one-half of outgrowth cultures were spread onto one 9-mm petri dish to obtain 300-500 independent colonies, and 192 random clones were screened using Sanger sequencing to create a collection of consolidated mutant strains covering each of 31 NNK sense codons. For SDM constructs, one-twentieth of 1 mL of the outgrowth cultures was spotted on LB-Kan agar in one well of 12 well plates to obtain 10-30 independent colonies, and screening 2 random clones was generally sufficient for identifying at least one correct construct.

To create the 4-site combinatorial mutants with Golden Gate assembly, we split the *rhlA* gene into four parts including 1-264 bp, 265-387 bp, 388-483 bp, and 484-888 bp, which encode 1-88 aa, 89-129 aa, 130-161 aa, and 162-296 aa of the RhlA protein (Figure S5 and Table S6). The split sites and linker sequences were designed using the NEB Golden Gate Assembly Tool (https://goldengate.neb.com/). Each of the four *rhlA* gene fragments was separately inserted into a receiver vector Lv0-ccdb via Golden Gate assembly using BsmBI, and the resulting WT substrate plasmid was then used as templates to generate the other 19 mutant substrate plasmids via Gibson assembly for SDM as abovementioned. All 80 substrate plasmids for 4 target residues were normalized to a final concentration of 30 ng/μL and stored in a 96-well storage plate. To construct a specific target sequence, four corresponding substrate plasmids were selected using a custom cherry-picking script of the liquid-handling robot (Figure S6) and then combined with the receiver plasmid pRSF-Duet-rhlb-ccdb for setting up Golden Gate assembly in 96-well PCR plates. The resulting reaction mixtures were used to transform chemically competent *E. coli* DH5α by heat shock, and 1/20 of 1 mL of the outgrowth cultures was spotted on LB-Kan agar in one well of 12-well plates to obtain 10-30 independent colonies. For each Golden Gate assembly reaction, screening of two random clones by Sanger Sequencing was generally sufficient to identify at least one correct construct.

#### Inducible production and detection of mono-RLs in E. coli

The plasmid DNA of SSM and combinatorial mutants isolated from DH5α was used to transform chemically competent *E. coli* BL21(DE3) to construct production strains. Transformants were obtained on LB-Kan agar in 24-well microplates after 20 h incubation at 37°C, and three independent clones were transferred into 2 mL of LB-Kan in 24-deep well plates. After 16 h cultivation at 37°C, 20 μL of stationary-phase cultures were used to inoculate 2 mL of fresh LB-Kan medium and incubated at 37°C for 3-4 h to reach the exponential growth phase (OD_600_=0.6-0.8). Then, IPTG was added to a final concentration of 1 mM for inducible mono-RL production at 30°C for 24 h (Figure S6).

For MALDI-ToF MS analysis (Figure S6), the resulting cultures were extracted with 2 mL of ethyl acetate with constant 600 rpm shaking for 0.5 h. Then, 1 μL of organic phase after 5 min centrifugation at 4,000 rpm was spotted onto MTP 384 AnchorChip steel targets preloaded with 0.3 μL of 2,5-dihydroxybenzoic acid (DHB) solution (10 mg/ml, acetoni-trile:H_2_O:trifluoroacetic acid (TFA) = 50:49.9:0.1, v/v). MALDI-ToF mass spectra were acquired using a Bruker Autoflex MALDI-ToF mass spectrometer (Bruker Daltonics) equipped with a frequency tripled Nd:YAG solid-state laser (λ=355 nm). Mass spectrometer calibration was performed using Peptide Calibration Standard Kit II (Bruker Daltonics). Spectral acquisition was performed in positive reflection mode with pulsed ion extraction and a mass range of 400-1,000 Da, and 1,000 shots (5 different positions into each spot and 200 shots per sub-spectrum) were accumulated. The laser footprint was set to “Ultra” at a ~100 μm diameter, and 1,000 laser shots were fired at 1,000 Hz. The spectra data files obtained from MALDI-ToF MS were imported into ClinProTools software (Bruker) and converted to CSV format for feature extraction normalized using total ion count. For mono-RL quantification, the intensities of [M+Na]^+^ ions of Rha-(C_8_-C_10_) (m/z 499.3) and Rha-C_10_-C_10_ (m/z 527.3) were applied (Figure 7b). The total mono-RL production was defined as Sum = Intensity_m/z 499.3 ion_ + Intensity_m/z 527.3 ion_, and relative Rha-(C_8_-C_10_) production was defined as Ratio = Intensity_m/z 499.3 ion_ / (Intensity_m/z 499.3 ion_ + Intensity_m/z 527.3 ion_). Then, normalized total mono-RL production relative to WT was defined as Sum_norm = Sum(mutant) / Sum (WT), and normalized relative Rha-(C_8_-C_10_) production was defined as Ratio_norm = Ratio (mutant) / Ratio (WT). For BO-EVO, the fitness values were defined as Fitness = Sum_norm × Ratio_norm (Table S4).

## RESULTS

### Design principles of BO-EVO and software implementation

BO-EVO integrates BO and evolutionary algorithms with complementary advantages. The computational time of BO increases exponentially with the number of targeted residues (N) when evaluating the whole protein combinatorial space (20^N^). To improve algorithm scalability, BO-EVO restricts the search to sub-spaces generated by random mutation of a parent sequence (evolution). The parent sequence is either sampled from measured sequences by fitness-based sampling or set as a newly proposed sequence depending on the confidence level of the surrogate model (Figure 1a). On the other hand, BO allows enhanced exploration than greedy evolutionary algorithms. We used GPR as surrogate models to quantify uncertainty. GPR is trained on measured sequences encoded by numerical protein sequence representations, such as ESM-1v (14). To prioritize mutant sequences for experiments, we utilized UCB as the acquisition function to negotiate exploration and exploitation. Once reaching a set budget for each round (i.e., 24, 96, 384 sequences), a mutant batch is experimentally created and analyzed. The resulting sequence-function data are used to refine the surrogate model for guiding further engineering rounds (Figure 1b).

**Figure 1.**
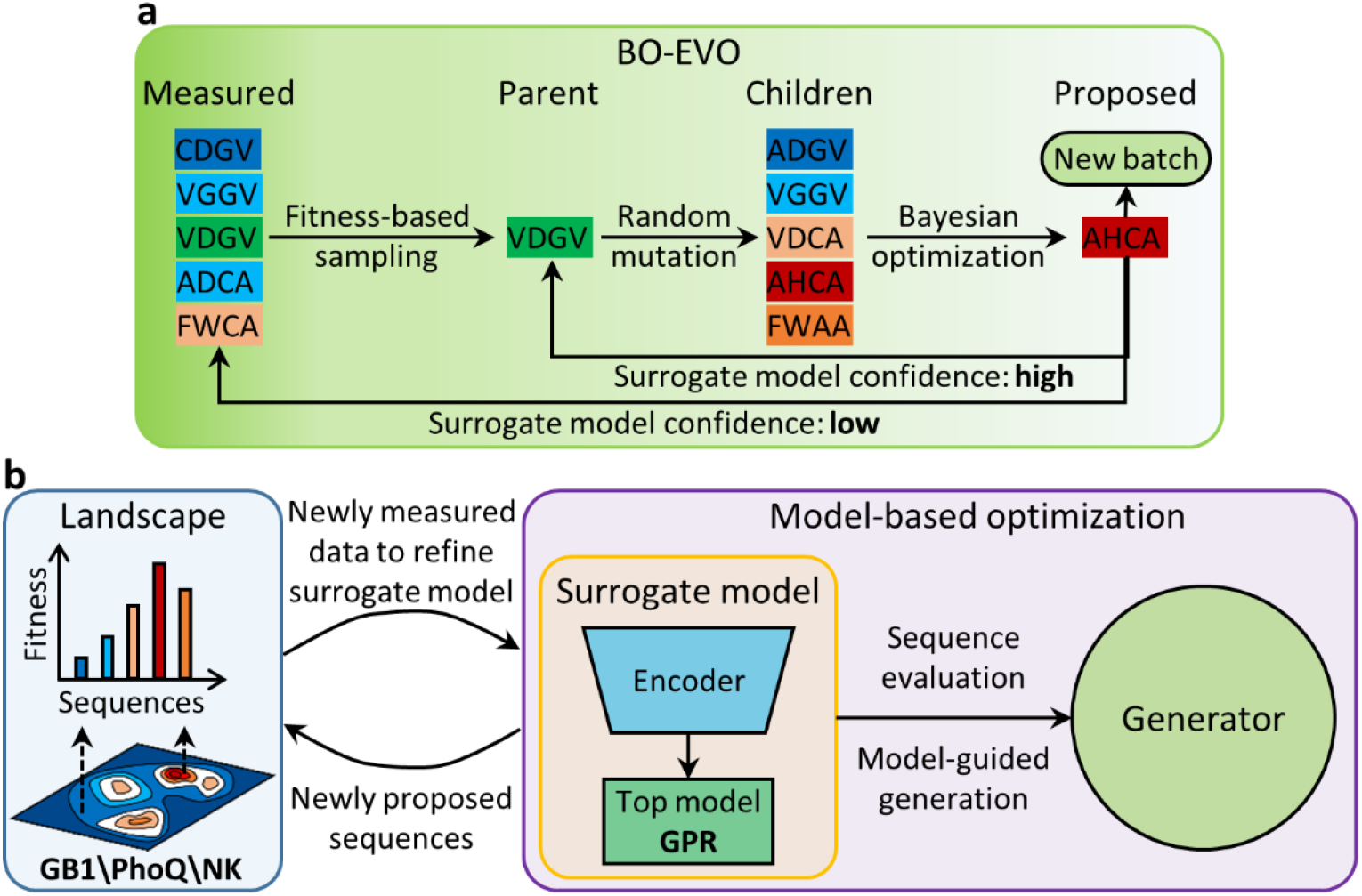
Scheme of BO-EVO. **a** Conceptual diagram of BO-EVO. Iteratively, candidate sequences (children) are generated by random mutation of a parent sequence, and one new sequence is proposed by BO on the candidates. **b** Software framework (FAST-HIT) consists of four modules: landscape, encoder, top model, and generator. The generator module proposes batch sequences by interacting with the other three modules.

To facilitate fine-tuning of BO-EVO design specifications, we implemented FAST-HIT as an in-house, model-based optimization software framework (Figure 1b). FAST-HIT consists of four modules, including landscape, encoder, top model, and generator. The landscape module provides sequence queries of different types, such as empirical landscapes like GB1 (35) and PhoQ (36), a mathematical landscape generated using the NK model (37), and an interactive landscape coupled with wet-lab measurement. The encoder module transforms amino acid sequences into numerical encoding. In addition to the categorical Onehot encoding, several learned representations and one physiochemical encoding are also included. The top model module executes training and evaluating diverse ML models. Here, we primarily implemented GPR model with GPyTorch (39) for accelerated model training and evaluation. The generator module proposes sequences through interaction with the other three modules.

### Moderate batch size is beneficial for low-round measurement

We first developed BO-EVO on an empirical fitness landscape of GB1 (35), which consists of the sequence-affinity data of a 4-site SSM library (20^4^=160,000 possible sequences). To run a simulation, 20 starting sequences were selected to be diverse in hamming distance to the global optima (fitness value set as 1.0) and with similar fitness levels as the wild-type (WT) sequence (fitness value ~ 0.1). For each of the 20 starting sequences, five random seeds were used to initiate BO-EVO, and hence in total 100 simulations were conducted when evaluating each algorithm design. Five metrics were used to evaluate algorithm performance. For round-wise evaluation, the cumulative maximum fitness and the success ratio to reach the global optima after each iteration were used; for the outcome, the maximum and mean fitness values of all the proposed sequences were used; and for model performance evaluation, the Pearson correlation coefficient was used.

To optimize the selection strategy for batch size and iteration number, which are key determinants for experimental budgets, two types of BO-EVO simulations were performed using the simple Onehot encoding. We first fixed the iteration number as 4, so that the protein engineering campaign can be completed in about one month. We observed that the maximum fitness increased with larger batch sizes, and the global optima can be consistently reached when the batch sizes were equal to or greater than 384, or the total sample budgets were equal to or greater than 1,536 (Figure 2a). Interestingly, the mean fitness of all proposed variants first increased and then decreased with greater batch sizes, possibly due to the limited number of high-fitness variants in the sequence space (only 2.31% variants exhibiting fitness higher than WT (35)), as well as the enhanced exploration in less-fitted regions with larger batches by BO-EVO.

**Figure 2.**
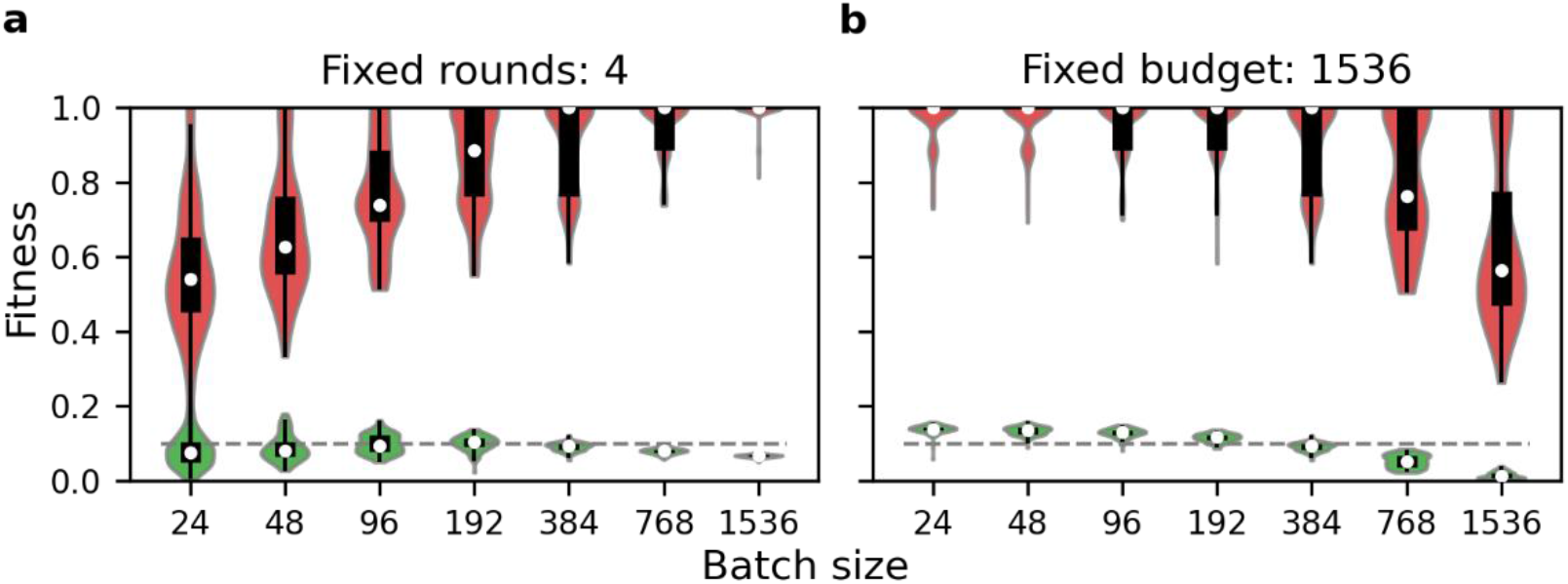
BO-EVO simulation on different experimental batch sizes. **a** Fixed iteration number (4). **b** Fixed total sample budget (1,536/160,000=0.096%). For each batch size, the maximum (red) and mean (green) fitness of all the proposed variants in 100 simulations were reported; the Violin plot integrates estimation of kernel density function; the box plot depicts the median, the first quantile, and the third quantile of fitness values. The gray dashed line plots the fitness value (0.1) of the WT sequence.

For the second type of simulation, we fixed the total sample budget as 1,536 (0.096% of the whole design space 160,000), which was the minimum sample number required to consistently reach the global optima within four BO-EVO iterations (Figure 2a). We noted that the maximum and mean fitness of 1,536 proposed mutants deteriorated with increasing batch sizes and decreasing iteration numbers (Figure 2b). These results indicated that algorithm-experiment feedback is necessary to improve BO-EVO performance on sequence design. Although favored in the second simulation, large iteration numbers are more time-consuming, as each experimental round generally takes a fixed turnabout of 5-7 days to execute. Overall, we decided to continue with moderate iteration numbers (4–5) and batch sizes (384) for BO-EVO experiments.

### Informative encodings and mutant data improve surrogate model performance

Protein sequences must be converted to numerical vectors as ML algorithm inputs. Here we evaluated a range of sequence encoding strategies, including simple categorical encoding (Onehot), physicochemical encoding (Georgiev), and learned representations, for their impact on surrogate model performance. In particular, Georgiev encoding is more informative than Onehot encoding by including a manually selected set of physicochemical parameters of each canonical amino acid (46). On the contrary, protein sequence featurization can be achieved through representation learning (53), assuming the physicochemical, structural, and evolutionary constraints may be discovered by observing naturally occurring protein sequences by unsupervised ML models. We examined two types of models, including UniRep (12) learned by a multiplicative recurrent neural network, as well as TAPE (13), ESM-1v (14), and ProtT5-XL-U50 (15) that are learned by transformer models (54) with different model architectures and training data sets (Table S3).

First, we evaluated the impact of encoding strategies on surrogate model quality, in terms of the Pearson correlation coefficient (Figure 3a). We first sampled a holdout test set of 384 samples from the GB1 landscape and then built a new empirical landscape with the remaining data. Simulations were conducted on this new empirical landscape and surrogate models were evaluated on the holdout test set. We found that model performance was continually improved with more rounds of iteration for all the encoding strategies, indicating model improvement was resulted from interactions with landscape data. On the other hand, informative encodings did not guarantee good models. Compared with the Onehot encoding, the use of learned encodings such as UniRep, ESM-1v, and ProtT5-XL-U50 was desirable, whereas the use of TAPE or Georgiev encodings was unfavoured (Figure 3a).

**Figure 3.**
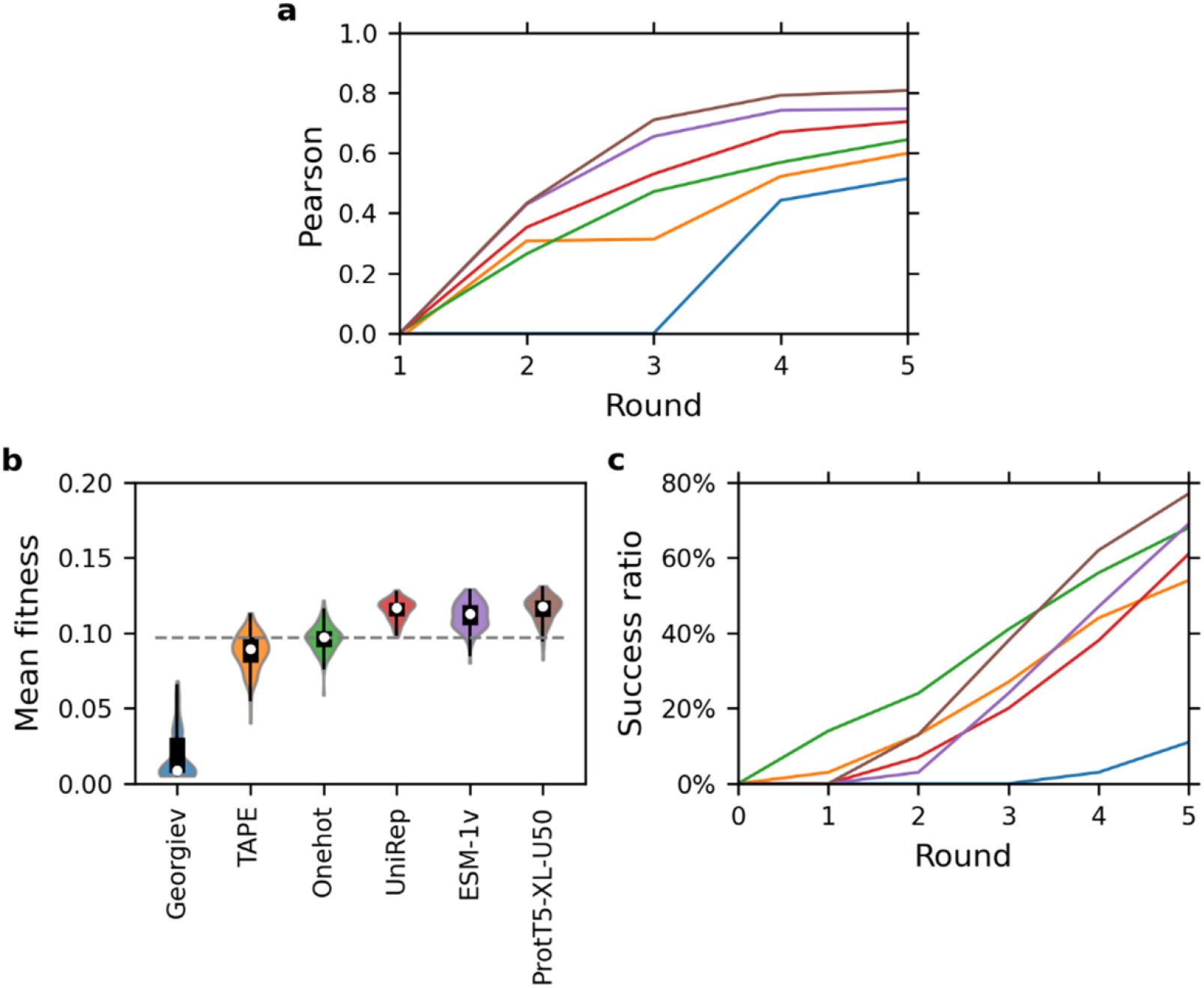
Comparison of protein sequence encoding strategies on BO-EVO performance. **a** Surrogate model performance at each round, evaluated with Pearson correlation coefficient on a holdout test set. **b** Mean fitness obtained in all the proposed variants for each encoding strategy. **c** Success ratio reached until each round. We use the violin plot as in Figure 2. The gray dashed line shows the median of mean fitness with Onehot encoding. All the subplots share the same color map.

Then, we examined how sequence encodings may affect BO-EVO performance. When examining the mean fitness of all the proposed sequences after the fifth iteration (Figure 3b), a similar trend of ProtT5-XL-U50 ~ ESM-1v ~ UniRep > Onehot > TAPE >> Georgiev encodings was observed as in Figure 3a, suggesting a strong correlation between the performance of the surrogate model and that of BO-EVO. When evaluating the success ratio after a given iteration (Figure 3c), however, the trend became a little different. In the first three rounds, the simple Onehot encoding outperformed all other strategies. After the fifth iteration, the performance rank was ProtT5-XL-U50 > ESM-1v ~ Onehot > UniRep > TAPE >> Georgiev, which again correlated well with the case for surrogate models (Figure 3a). Overall, ProtT5-XL-U50 and ESM-1v strategies were the top choices. For subsequent investigation, we proceeded with the ESM-1v encoding due to its smaller model size and higher computational efficiency compared with ProtT5-XL-U50. Notably, the observation that more complex encodings performed only marginally better at best than the simple Onehot encoding agrees with other recent studies (17, 55, 56).

Because landscape information enhanced surrogate model quality (Figure 3a) and hence BO-EVO performance (Figures 3c), we further explored whether single-residue SSM data, which are easy to measure and small in scale (20×N, e.g., N=4 for the GB1 landscape), can be beneficial when used for model initialization (“warm start”). Relative to the cool-start strategy that only fed one sequence-fitness data to surrogate models, the warm-start strategy resulted in a substantial gain in the success ratio (~10%) throughout the different rounds of simulated iterations (Figure 4). The warm-start strategy was therefore recommended for BO-EVO implementation.

**Figure 4.**
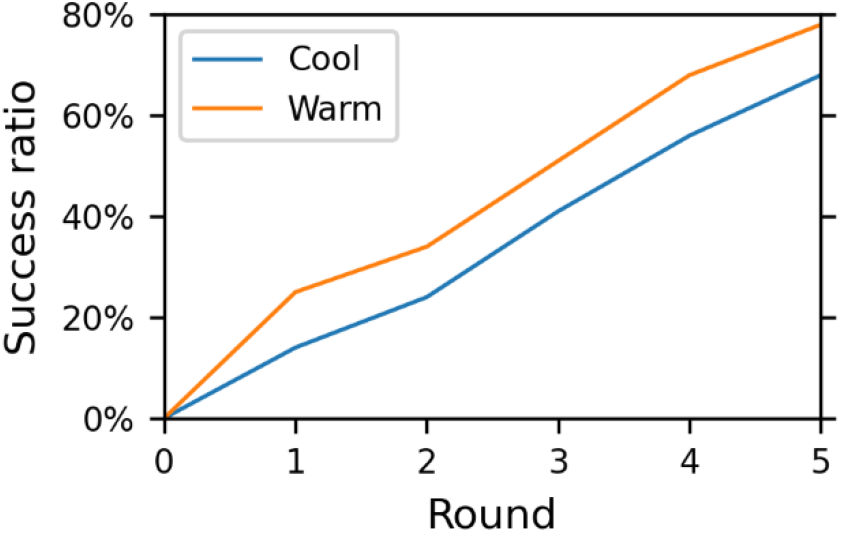
A warm start enhanced BO-EVO success ratios. In particular, after the fifth round, BO-EVO reached the success ratio of 78% and 68% for the warm-start and cool-start strategies, respectively.

### Comparison of fitness exploration algorithms

Next, we benchmarked BO-EVO with pure evolutionary (AdaLead (44)) and pure BO algorithms to examine the necessity to combine these two exploration strategies; and random mutagenesis (Random) was also evaluated as a baseline (see Table S2 for algorithm setup details). The performance of Random was the worst of the four algorithms as expected (Figure 5). In terms of the round-wise success ratios (Figure 5a) and the maximum and mean fitness of all the proposed sequences after five rounds (Figure 5b), AdaLead was better than Random by greedy exploitation on top variants with high predicted fitness, but this pure evolutionary algorithm was substantially worse than BO-EVO and BO, both of which utilized the UCB acquisition function. These results demonstrated the importance to consider both variant fitness and model uncertainty when exploring a rugged fitness landscape. On the other hand, by exhaustive exploration of the whole design space (160,000 sequences) in each iteration, pure BO achieved better performance than BO-EVO (Figure 5), which only evaluated 3,072 sequences (1.92% of the whole design space) *in silico* per round. Although the performance of BO-EVO was not as good as pure BO, the computational time of BO-EVO was almost constant for exploring combinatorial mutagenesis landscapes, whereas the computational time of pure BO scaled exponentially with targeted residue numbers and rapidly became intractable (Figure S4).

**Figure 5.**
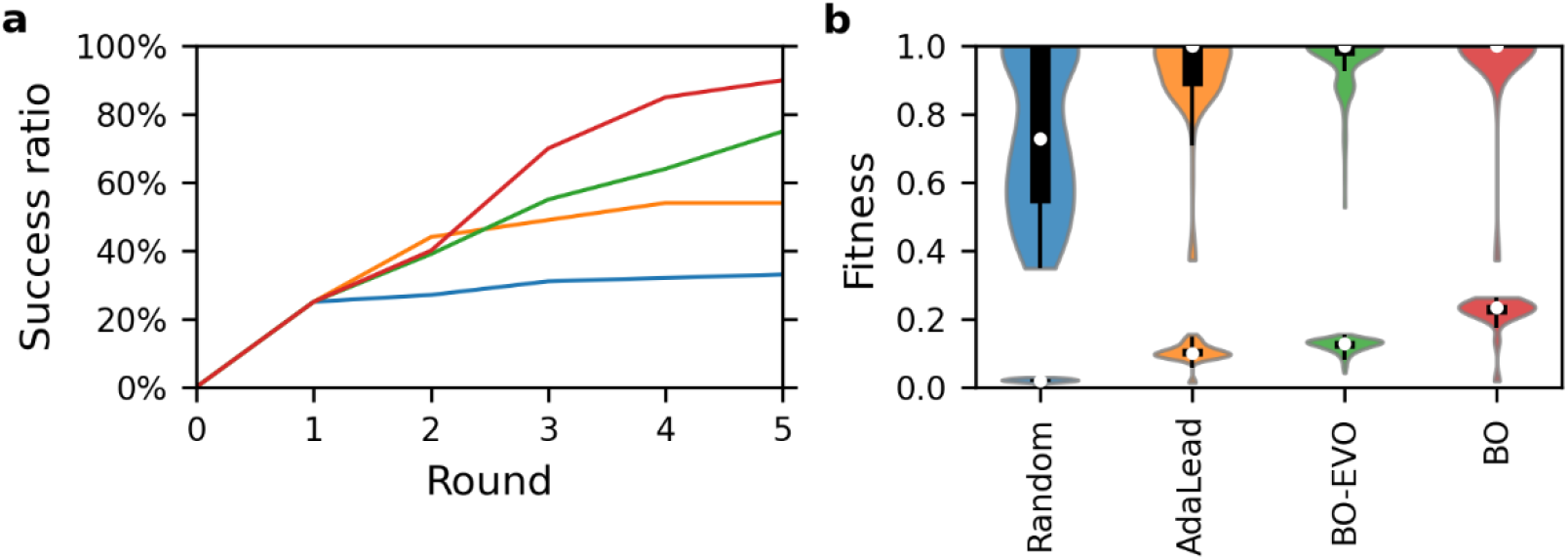
Fitness exploration algorithms. **a** Success ratio reached until each round. **b** Maximum (on the top) and mean (on the bottom) fitness obtained in all the proposed variants. The same violin plot is used as in Figure 2. All the subplots share the same color map.

### The ruggedness of fitness landscapes challenges the performance of BO-EVO

We further assessed the general applicability of BO-EVO in exploring multi-site combinatorial protein libraries using two additional fitness landscapes: one is empirical, and the other is simulated. The empirical fitness landscape reflects the protein-protein interaction (PPI) between *Escherichia coli* protein kinase PhoQ mutants and its substrate PhoP (36). To the best of our knowledge, the GB1 and PhoQ fitness landscapes are the only two comprehensive data sets on 4-site combinatorial libraries in literature. The simulated fitness landscapes were generated based on the NK model (37), where the parameter N denotes protein sequence length and K denotes a tuneable degree of epistasis.

To better mimic empirical fitness landscapes, the original NK model (37) was reformulated in three subsequent steps. First, the fitness distribution of empirical landscapes often obeys exponential distribution (3). Accordingly, we independently sampled the “fitness table” of a select residue from exponential distribution instead of uniform distribution. Second, only a small number of key residues exert substantial impacts on a target function. Therefore, we proposed a weighted average of the “fitness contribution” of each residue sampled from the Zipf distribution. Third, because functional proteins only constitute a tiny fraction of the whole sequence space (2), we simply set the low-fitness variants to “non-functional” with a cut-off threshold and created a black hole-filled, exponentially distributed NK landscape (Figure S1). The final reformulated NK landscape was applied for BO-EVO simulation with N=4 for 4-site combinatorial landscapes and various degrees of epistasis (K=0, 1, 2, 3).

Various measures of epistasis in the literature appear to be equivalent (42), and here we used the ruggedness-to-slope ratio (43), *r/s*, to measure landscape ruggedness. For NK landscapes, the landscape ruggedness scaled exponentially with K (Figure 6a). The ruggedness of the GB1 and PhoQ landscapes were comparable to the K=0 landscape and K=1 landscape, respectively (Figure 6b, purple star and blue triangle), indicating that the fitness landscape of PhoQ exhibited stronger epistasis than that of GB1. For exploring the simulated NK landscape using BO-EVO, the success ratio of reaching the global maximum decreased logarithmically with ruggedness (Figure 6b, green dots). On the other hand, although the empirical GB1 landscape and the simulated K=0 landscape had similar levels of epistasis, the former (Figure 6b, purple star) proved a more difficult task for BO-EVO than the latter (Figure 6b, green dot). Furthermore, BO-EVO achieved similar success ratios for the GB1 and PhoQ landscapes (Figure 6b), whereas the fitness landscape of PhoQ was more rugged than that of GB1. These results indicated that BO-EVO generalized reasonably well for fitness landscapes with moderate epistasis, and landscape ruggedness is one major challenge to BO-EVO to locate the global optima.

**Figure 6.**
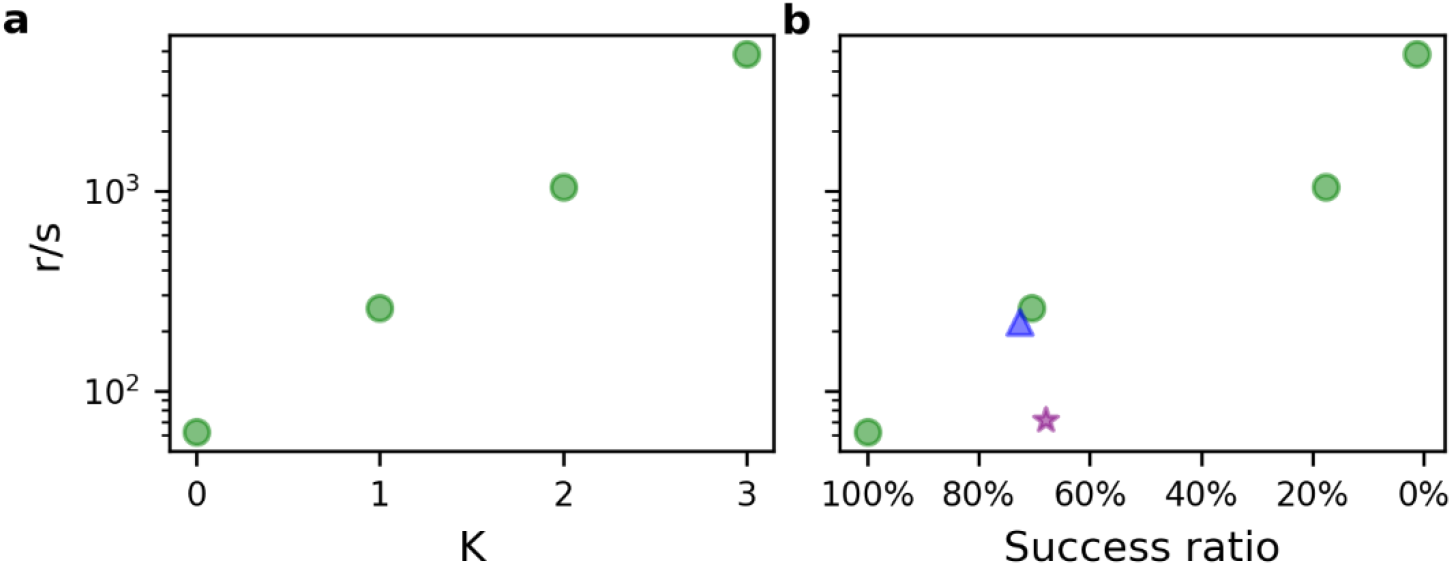
Generalization of BO-EVO to empirical and simulated landscapes with diverse ruggedness. **a** Ruggedness of NK landscapes. **b** Success ratio after the fifth BO-EVO round. The results for the landscapes of NK, GB1, and PhoQ are shown with green dots, a purple star, and a blue triangle, respectively.

**Figure 7.**
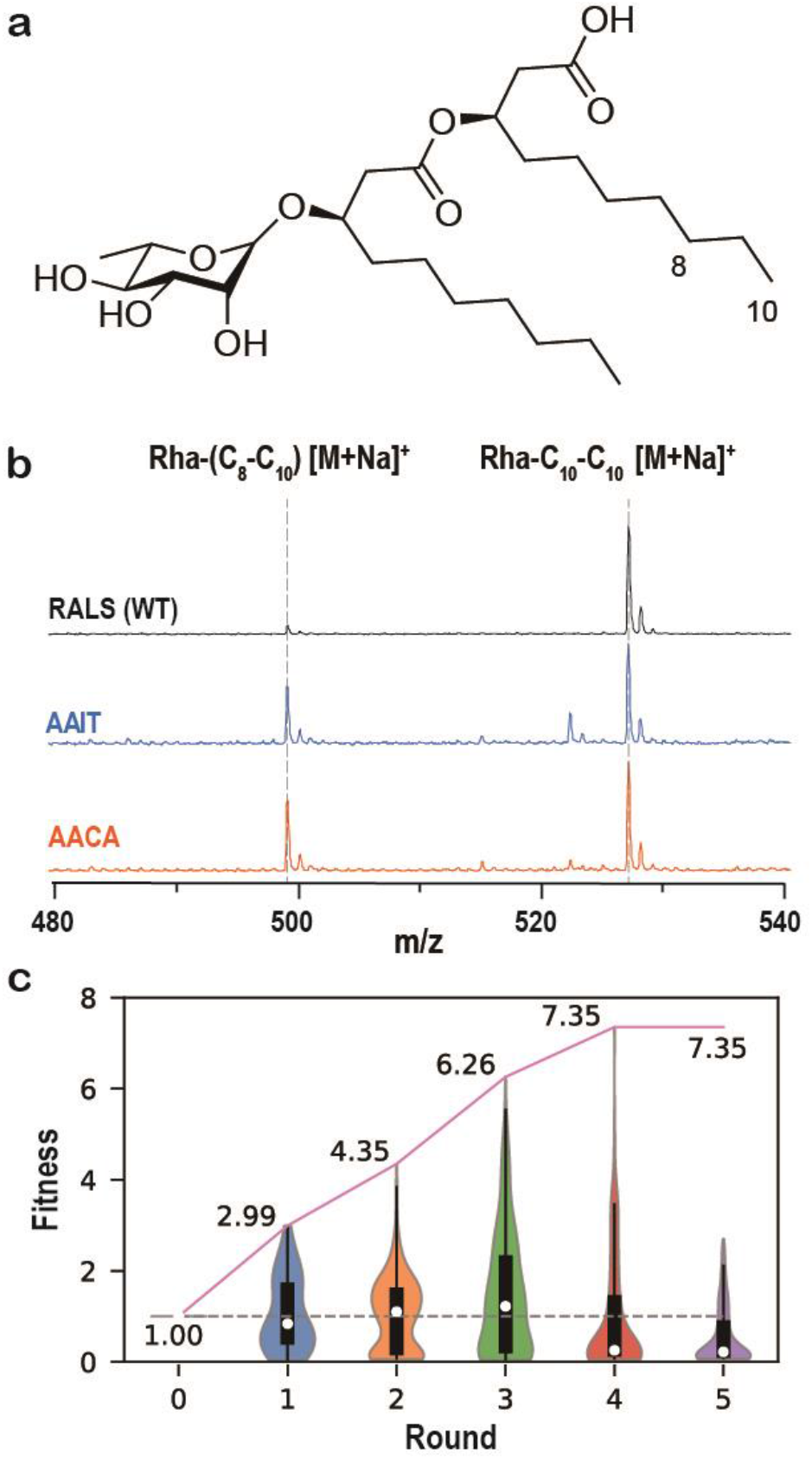
BO-EVO guided 4-residue combinatorial SSM engineering of RhlA. **a** Molecular structure of mono-RL Rha-C10-C10. **b** MALDI mass spectra of mono-RLs produced from WT and representative RhlA mutants (Table S4) with amino acid compositions at Residues 74, 101, 148, and 173 labelled. **c** Normalized production of Rha-(C8-C10) relative to WT (set as 1, dashed line) was reported. The cumulative maximum fitness was numerically labelled and plotted as a solid line. For each round, the same violin plot was used as in Figure 2.

### Enzyme engineering via algorithm-experiment iterations by BO-EVO

Next, we sought to apply BO-EVO to a real-world protein engineering task via iterative feedback between ML models and robotic experiments. RhlA is a key enzyme responsible for synthesizing the lipid moieties of rhamnolipids (RLs), an important biosurfactant. The enzyme specificity of RhlA determines the chemical structures of the lipid moieties (Figure 7a), which further affect the physiochemical and biological activities of the corresponding RL molecules. However, it proves difficult to alter the enzyme specificity of RhlA through either (semi-)rational design or directed evolution (57, 58). To apply BO-EVO for RhlA engineering, we developed a robotic protocol to build and test any 384 out of the 160,000 members in the 4-residue combinatorial SSM libraries of RhlA (Figures S5). Using this protocol (Figure S6), which will be reported in a separate work, each experimental round took less than one week to complete. To measure enzyme specificity as the target function, we applied a previously reported matrix-assisted laser desorption/ionization time-of-flight (MALDI-ToF) mass spectrometry (MS) assay (23, 59) to quantify two RL products Rha-(C_8_-C_10_) and Rha-C_10_-C_10_ (Figure 7b), in the liquid cultures of recombinant *Escherichia coli* as the production host. In microplate cultures, *E. coli* cells harbouring the WT RhlA produced Rha-C_10_-C_10_ as the main product (Figure 7b), and we aimed at shifting the product specificity towards smaller Rha-C18 product in RhlA mutants. The normalized production levels of Rha-(C_8_-C_10_) relative to WT were used to measure fitness, where the fitness of the WT sequence was set to 1.

To apply BO-EVO, we selected Arg74, Ala101, Leu148, and Ser173 (RALS) as the four target residues for combinatorial mutagenesis, because many mutations at these ligand-binding residues substantially enhanced Rha-(C_8_-C_10_) production. The single-residue SSM data of these 4 residues were used for a “warm start” (Round 1), which was desirable for improving outcomes (Figure 4). For BO-EVO iterations, we observed a round-wise increase in the cumulative maximum fitness, which reached 7.35 (AACA sequence) as the highest value in Round 4. For secondary confirmation, the top 5 hits from Round 4 were subjected to plasmid re-transformation and repeated MALDI-ToF MS analysis. As a result, although improvement in Rha-(C_8_-C_10_) percentiles was consistent with the first-pass screening, we observed substantial changes in the overall RL production levels (Table S4), possibly due to the limited quantitation capability of MALDI-ToF MS for mono-RL (59). The use of alternative high-throughput MS modalities, such as the RapidFire ESI-MS system, may assist in reducing the discrepancy between primary and secondary screening in the future (60). Nonetheless, we successfully identified one RhlA mutant conferring a 4.8-fold improvement in RL-(C_8_-C_10_) production relative to WT (Table S4 and Figure 7c).

For comparison, we also performed simulated engineering iterations to explore the fitness landscapes of GB1 and PhoQ. Starting from the WT sequences, 10 BO-EVO rounds were executed for each simulation, and simulations were repeated 5 times for each protein (Figure S7). The patterns of round-wise changes were similar between RhlA (Figure 7) and GB1 (Figure S7a), exhibiting an initial increase and a subsequent decrease in the median and top fitness. For GB1 simulations, the turning point was near the round that identified the global optima. On the other hand, PhoQ proved to be an easier engineering target than RhlA and GB1 for BO-EVO, which identified the best-performing variant within 2 rounds in 4 out of 5 simulations (Figure S7b). Moreover, PhoQ simulations showed no obvious changes in the round-wise median and overall fitness, which were different from RhlA and GB1. The distinct performances of BO-EVO between PhoQ and GB1 may result from the different features of these two landscapes, such as WT fitness values, fitness distributions, and landscape ruggedness (Figure S1 & Figure S2).

## DISCUSSION

In this work, we developed a scalable and batched BO algorithm, BO-EVO, for guiding multiple rounds of robotic experiments to explore combinatorial protein fitness landscapes. For 4-site combinatorial mutagenesis, the total experimental budget of BO-EVO was reduced to 1,536 mutants or less than 1% of the theoretical library size of 160,000, where 4 iterations of web-lab measurement, model refinement, and mutant design were executed with a moderate batch size of 384. Within these sample budgets, BO-EVO achieved decent success rates (75%) to reach global optima on a range of simulated and empirical protein fitness landscapes. More importantly, a real protein engineering task was performed to modify the product specificity of RhlA, and substantially improved mutants were rapidly designed by BO-EVO within a month. To the best of our knowledge, this is the first report of algorithm-guided, automated protein engineering via multiple feedback rounds between ML models and robotic experiments.

Previously, a BO algorithm-guided pathway engineering platform, BioAutomata (27), was reported to robotically construct and evaluate 136 pathway variants out of all 13,824 possible library members, achieving a 1.77-fold higher titer in lycopene. While both BO-EVO and BioAutomata are based on similar biofoundry setups, the problem dimension of BO-EVO in this study is one magnitude higher than BioAutomata, urging advances in both algorithms and robotics. For the former, an evolutionary algorithm was applied to restrict the search space of BO for better computational efficiency; for the latter, new robotic protocols for synthetic biology and MS analysis were developed to create and profile a batch of 384 enzyme mutants within a week for rapid experiment-model feedback.

Several adaptive experimental design approaches, such as MLDE (17), cluster learning-assisted directed evolution (CLADE) (61), and ODBO (62), have utilized ML models for sampling-efficient protein engineering. There are two key differences between these approaches and our method. First, BO-EVO does not require prior knowledge of the structure or homolog sequences of a target protein, which may be unavailable or scarce. On the contrary, information derived from protein structures (e.g., ΔΔG) and homolog sequences were utilized by MLDE (17) and CLADE (61), respectively, as zero-shot predictors to exclude possible zero- or low-fitness mutants from experimental sets. Second, BO-EVO does not need to evaluate the whole design space *in silico* by restricting sequence design via evolutionary algorithms. In particular, 3,072 candidates (1.92% of all 160,000 possible sequences for 4-site combinatorial mutagenesis) were evaluated by surrogate models during each BO-EVO iteration. Contrariwise, MLDE, CLADE, and ODBO employed fitness models for *in silico* evaluation of all possible sequences. On top of fitness prediction, CLADE applied deep hierarchical clustering of the whole design space based on physicochemical sequence encodings, and ODBO utilized an outlier mining algorithm to identify low-fitness candidates from all possible sequences. However, the computational time of full-space prediction will increase exponentially and soon become intractable with high-dimensional problems (Figure S4). With the ever-decreasing experimental time and cost required for robotic sequence-function profiling, the sampling efficiencies of both experiment and computation need to be carefully balanced.

In the future, BO-EVO can be further improved in uncertainty quantification, mutant residue selection, and experimental capabilities. For uncertainty quantification, more accurate and computationally tractable models, including ensemble methods, Bayesian neural network (63), and evidential deep learning (64), can be applied to replace GPR. Furthermore, the four target residues of RhlA were manually picked in this study based on previous experiments, and it is desirable to apply algorithms to guide mutant residue selection (65). To this end, MutCompute (66) has trained a self-supervised convolutional neural network (CNN) on protein structure data to identify hot spots in PETases for introducing stabilizing mutations (67). Third, we studied a 4-site combinatorial library with a theoretical size of 160,000 by creating and screening 384 protein mutants per week. Further improvement in robotics may assist in exploring more complex variant libraries by BO-EVO. It may also be interesting to comprehensively profile every RhlA mutant in this particular library with enhanced experimental throughput so that the performance of BO-EVO can be examined in retrospect. Overall, we successfully developed a new approach for protein engineering via algorithm-guided robotic experiments for combinatorial mutagenesis.

## DATA AVAILABILITY

All data generated and analyzed in this study are included in the manuscript and supplementary information.

## ACKNOWLEDGEMENT

We thank Professor Lei Dai for the helpful discussion on NK landscapes. We thank the Shenzhen Synthetic Biology Infrastructure for robotic workflow development.

